# Scalable and efficient single-cell DNA methylation sequencing by combinatorial indexing

**DOI:** 10.1101/157230

**Authors:** Ryan M. Mulqueen, Dmitry Pokholok, Steve Norberg, Andrew J. Fields, Duanchen Sun, Kristof A. Torkenczy, Jay Shendure, Cole Trapnell, Brian J. O’Roak, Zheng Xia, Frank J. Steemers, Andrew C. Adey

**Affiliations:** Department of Molecular & Medical Genetics, Oregon Health & Science University, Portland, Oregon, USA; Advanced Research Group, Illumina, Inc., San Diego, California, USA; Department of Molecular Microbiology & Immunology, Oregon Health & Science University, Portland, Oregon, USA; Computational Biology Program, Oregon Health & Science University, Portland, Oregon, USA; Department of Genome Sciences, University of Washington, Seattle, Washington, USA; Howard Hughes Medical Institute, Seattle, Washington, USA; Knight Cardiovascular Institute, Portland, Oregon, USA

## Abstract

Here we present a novel method: single-cell combinatorial indexing for methylation analysis (sci-MET), which is the first highly scalable assay for whole genome methylation profiling of single cells. We use sci-MET to produce 2,697 total single-cell bisulfite sequencing libraries and achieve read alignment rates of 69 ± 7%, comparable to those of bulk cell methods. As a proof of concept, we applied sci-MET to successfully deconvolve the cellular identity of a mixture of three human cell lines.

## Main

The fundamental challenge of identifying and characterizing the molecular properties of every cell type in the body has recently entered the realm of possibility^1^. High-throughput singlecell transcriptome^2–4^ and chromatin accessibility profiling assays^5–7^ have dramatically improved our ability to uncover latent cell types within complex tissues and dynamic cell states during differentiation. These same possibilities extend to DNA methylation; a covalent addition of a methyl group to cytosine bases in the genome that largely serves in a repressive role^8^. DNA methylation occurs at a high rate in cytosine-guanine dinucleotides (CG), with cytosine methylation in non-CG sites (CH) occurring rarely and only in select tissues^9^. Both CG and CH methylation have cell type-specificity and are the subject of active modification in developing tissues^8^. DNA methylation can be probed at base-pair resolution using the deaminating chemistry of sodium bisulfite treatment before or after the generation of sequencing libraries ***(e.g.*** as in whole genome bisulfite sequencing, WGBS)^10,11^. Despite the comprehensive nature of these assays, key aspects of methylation architecture and dynamics remain elusive, with cell type heterogeneity at the forefront of this challenge.

Recent work has optimized bisulfite sequencing to decrease input requirements to the single cell level (scWGBS)^12–15^. These assays have provided unique insights into environmental effects on methylome dynamics^13^, have uncovered new cell-types that form during hematopoesis^14^, and been coupled with scRNA-seq to directly study the relationship of DNA methylation to transcription^15^. However, these methods use a parallelized library generation strategy in which the WGBS protocol must be carried out on each cell in its own reaction vessel and thus remain low-throughput. Existing platforms to assess the transcriptome of single cells in high-throughput largely rely on droplet-based microfluidics strategies by sequestering single cells into individual droplets along with a substrate harboring barcoded oligonucleotides for cell-level indexing^2,16^. This approach is not readily adaptable to genome-wide DNA methylation profiling due to harsh bisulfite conversion chemistry, which destroys cell and nuclear integrity (preventing conversion prior to droplet encapsulation), and cannot be carried out in a single reaction buffer (preventing both conversion and barcoding in the same droplet). Furthermore, alignment rates for traditional scWGBS libraries are on the order of 25 ± 20%, much lower than for the equivalent bulk protocol^12–15^, which markedly increases the cost of obtaining sufficient aligned read counts.

Recently, we described a platform for combinatorial indexing to acquire long-range sequence information^17^ that has been extended to single cell assays for open chromatin (sci-ATAC-seq, scTHS-seq)^5,7^, chromosome conformation (sci-Hi-C)^18^, RNA transcript abundance (sci-RNA-seq)^19^, and whole genome sequencing (sci-DNA-seq)^20^. The key to this combinatorial indexing platform is a two-tier indexing strategy where DNA (or RNA) within nuclei (or cells) is modified with a barcoded adaptor corresponding to one of 96 (or 384) wells while maintaining nuclear integrity. All reactions are then pooled, followed by the redistribution of a limited number of these pre-barcoded nuclei into each of a new set of wells such that the probability of two nuclei harboring the same initial barcode ending up in the same well is low. PCR is then used to incorporate a second index (96 or 384) with corresponding sequencing adaptors. Thus, each library molecule contains two indexes: one from the initial barcoding, and the second from PCR, allowing single-cell discrimination. For sci-DNA-seq^20^, we deployed a nucleosome depletion strategy prior to combinatorial indexing which enabled relatively uniform coverage of the genome, a key component for extending this concept to measure genome-wide DNA methylation in single cells.

We adapted our combinatorial indexing strategy to single cell WGBS (Fig. 1a). In contrast to previous work^11^, transposomes are loaded with C-depleted oligonucleotides and thus unaffected by bisulfite treatment. This approach reduced synthesis costs by avoiding methylated adaptors and improved downstream PCR amplification while maintaining library size. Due to low GC content, the C-depleted adapters possess lower melting temperatures than standard sequencing primers. To overcome this, we utilized sequencing primers modified with locked nucleic acids (LNA) to mimic standard melting temperatures. The second (5’) adaptor is then incorporated after pooling, redistribution, bisulfite conversion, and cleanup by performing one or more rounds of random primer extension, as in traditional scWGBS protocols^12^.

**Figure 1.**
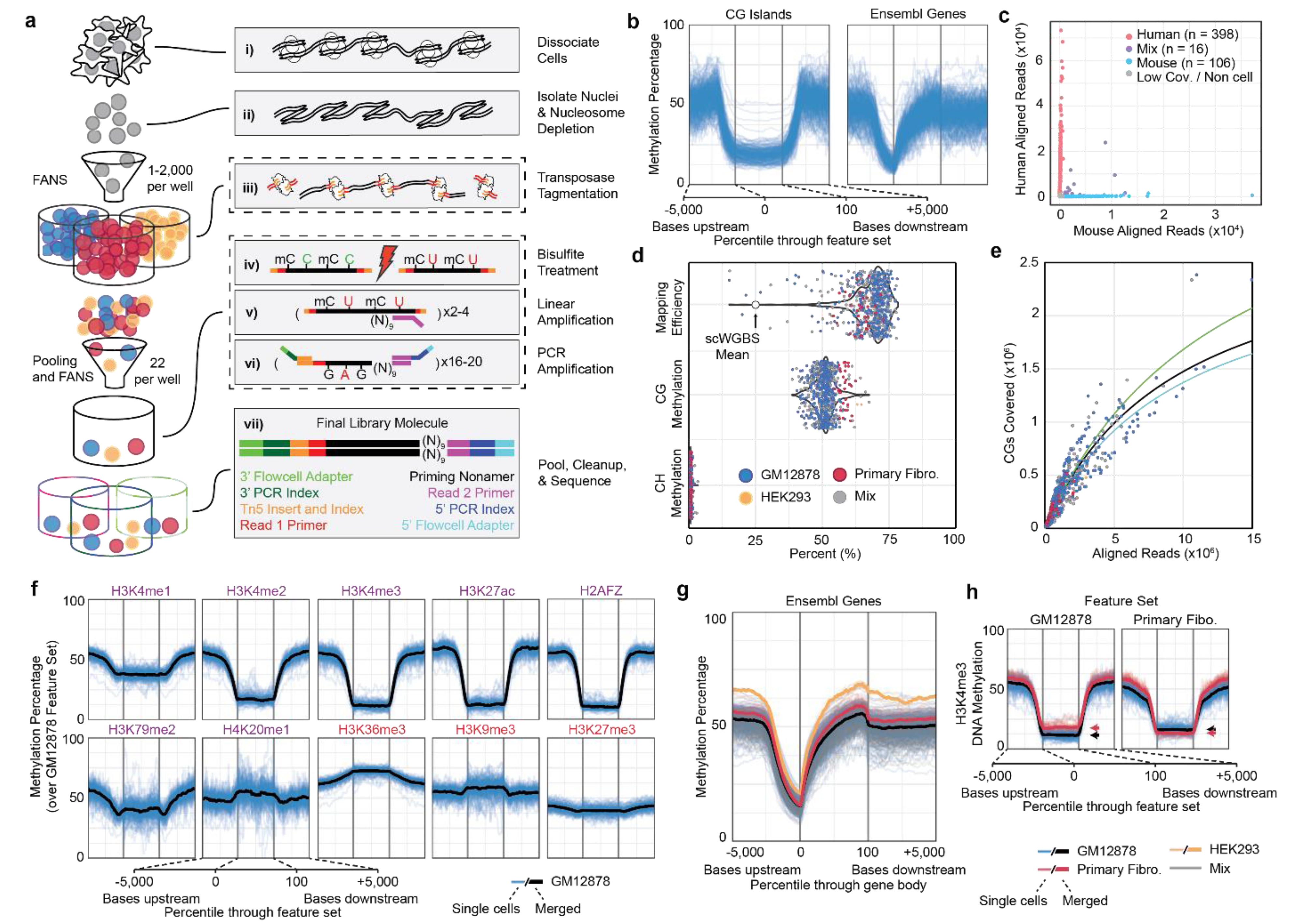
Sci-MET assay and performance. **a.** The sci-MET workflow. i) cells are obtained and contain nucleosome-bound DNA, ii) nuclei are then isolated and subjected to nucleosome depletion, iii) 1-2,000 nuclei are deposited into each well of a 96-well plate using fluorescence activated nuclei sorting (FANS) and reacted with transposase complexes with indexed, C-depleted, 3’ adaptors, iv) all wells are pooled from the transposase reaction and 22 nuclei are deposited into each of a set of new wells such that the probability of two nuclei sharing the same transposase index is low, several wells have unmethylated lambda genomic DNA that has been pre-transposed with a unique index spiked in, and then all wells are subjected to bisulfite treatment, v) after bead-based cleanup, two to four rounds of linear amplification is performed using a 5’ adaptor primer that terminates in nine N bases at its 3’ end, vi) PCR is then carried out using primers containing indexes corresponding to each well, vii) wells are pooled for sequencing. The final library molecule contains indexes that correspond to the transposase well and the bisulfite treatment + PCR well, which facilitates single-cell discrimination. **b.**Methylation rates for single GM12878 cells over CG islands (left) and gene bodies (right). 5,000 basepairs upstream and downstream of peaks are shown along with a percentile progression through the body of the features. As expected a marked decrease in methylation is observed in CG islands and in the bases leading up to transcription start sites. **c.**Human and mouse cells were mixed and carried through the sci-MET protocol using crosslinking and SDS nucleosome depletion to estimate barcode collision rates. **d.**Mapping efficiency (top), global CG methylation (middle) and global CH methylation (bottom) for a sci-MET preparation of a mix of human cell lines. Each point is a single cell colored by its identity. Alignment rates approach that of bulk bisulfite sequencing experiments, the mean using existing scWGBS technology is shown as a white point. **e.**The number of CG dinucleotides covered (in millions) by the total aligned reads per cell (in millions). Solid lines represent saturation curves for 3 rounds of linear amplification (blue), 4 rounds (green), or all cells (black). **f.**Methylation rates for GM12878 cells (blue lines, merged in black) over ENCODE ChIP-seq features captures the expected methylation profile. Purple feature names are for generally activating marks, red names are for generally repressive marks. **g.**Methylation rates for single cells (transparent lines) and merged for the three cell types (solid lines) over annotated genes. **h.**Methylation rates over GM12878 (left) and Primary Fibroblast (right) ENCODE H3K4me3 ChIP-seq peaks. GM12878 and Primary Fibro. cells are shown. Arrows indicate the mean for the entire length of the features set. Each respective cell type has a greater decrease in DNA methylation over the matching H3K4me3 peak set.

First, we assessed the viability of our strategy on a well characterized B-lymphoblast cell line (GM12878). From a single experiment (96-well transposase barcodes by 96-well PCR preparation; 96 × 96), we generated a library in which we could identify barcode combinations corresponding to 774 single cells. Sequencing this library to a low depth (mean 55,129 unique reads per cell) produced methylation profiles that closely matched expectation for the GM12878 cell line, including local depletion of methylation at gene promoters and CG islands (Fig. 1b, Supplementary Fig. 1,2). To assess the barcode collision rate (i.e. two nuclei of the same transposase barcode ending up in the same PCR well), we carried out sci-MET on a mix of human and mouse cell lines in a 96 × 40 experiment using lithium-based nucleosome depletion to characterize DNA methylation in 711 single cells (Supplementary Fig. 3). We classified collisions as barcodes with fewer than 90% of aligned reads to either the mouse or the human reference. Using this criteria, and assuming the same collision rate for undetectable human-human and mouse-mouse collisions, we estimate a total collision rate of 22%. This high rate is likely driven by decreased nuclear integrity during nucleosome depletion, so we repeated the experiment using an alternative crosslinking and SDS nucleosome depletion method. In a 96 × 32 experiment, we produced 520 single cell libraries with an estimated collision rate of 6% (Fig. 1c). This is near the expected rate from previous sci-methods. As with other combinatorial indexing strategies, the collision rate of sci-MET can be tuned by adjusting the number of nuclei sorted into each PCR well or number of barcodes.

We next sought to demonstrate the ability of sci-MET to deconvolve a mixed population of human cell types by profiling pure populations and a synthetic mixture of a B-lymphoblast (GM12878), primary inguinal fibroblast (Primary Fibro., GM05756), and embryonic kidney (HEK293) cell lines (Supplementary Fig. 1,2). In a 96 × 40 experiment using crosslinking and SDS nucleosome depletion, we characterized genome-wide methylation in 691 single cells passing QC filters, with a mean unique read count of 560,613 per cell; 107 cells had over one million uniquely aligned reads. Remarkably, we were able to achieve a mean alignment rate of 69 ± 7% (Fig. 1d), which is comparable to bulk WGBS rates and a substantial improvement over standard scWGBS (25 ± 20%)^12–15^. We suspect that this is due to the high efficiency of transposase-based adaptor incorporation and/or that single cells are not processed individually but in multiples, thus reducing adaptor dimers^11,21^. Given that the most substantial cost associated with single-cell methylation experiments is the ample raw sequencing that is required, our dramatically improved alignment rate has the potential to reduce associated sequencing costs by as much as 5 to 7 fold. Libraries also exhibited CG and non-CG methylation rates of 51 ± 4.0% and 0.77 ± 0.23% respectively (Fig. 1d), consistent with expectations and a high efficiency of bisulfite conversion (99.14% from lambda unmethylated control spike-in). We also observed increased unique CG coverage by unique read alignment (Fig. 1e), comparable coverage over annotated regions of the genome, with only a slight bias toward regions of open chromatin over regions with repressive marks (Supplementary Fig. 4), and increased unique CG nucleotide coverage when aggregating multiple cells - with 61 low-coverage cells required to cover approximately half of the CG dinucleotides in the haploid genome at our sequencing depth (Supplementary Fig. 5), all suggestive of a sufficient level of coverage uniformity for comprehensive methylome assessment. Furthermore, the methylation rate over functionally annotated regions conformed to expectations with a marked decrease in methylation at activating marks *(e.g.* H3K27ac) and an increase at repressed regions *(e.g.* H3K36me3; Fig. 1f). We also observed the expected decrease in methylation in the promoter region of genes for all cell types (Fig. 1g), and a greater magnitude of methylation rate change for cells matching the ChIP-seq sample (Fig. 1h, Supplementary Fig. 6).

Summarized methylation status^12^ was calculated for each cell across autosomal loci of the Ensembl Regulatory Build^22^, which contains regions of the genome known for transcription factor binding and epigenetic marks (Fig. 2b). We then performed Non-negative Matrix Factorization followed by t-distributed Stochastic Neighbor Embedding (NMF-tSNE) to project cells in two dimensional space (Fig. 2c). Distinct domains containing each respective cell type along with cells from the mixed population were observed. Cluster purity was further confirmed by the proportion of reads aligning to the Y-chromosome (specific to fibroblasts), as well as minimal bias pertaining to unique read count or global CG methylation percentage (Supplementary Fig. 7). Aggregating cells by cluster identity, we observed 47.7x10^6^ unique CG sites covered for the GM12878 cluster (599 cells), 5.35x10^6^ for the Primary Fibro. cluster (32 cells), and 3.31x10^6^ for the HEK293 cluster (10 cells). We next correlated the methylation rates with publically available WGBS datasets^23,24^ for the top 1,000 most variable regulatory regions. For each merged cluster, the two most highly correlated samples were of the same cell type, or the most similar cell line in the case of HEK293 (Fig. 2d, Supplementary Fig. 8). Hierarchical clustering on the Pearson correlation coefficients placed the HEK293 cluster in a clade with other aberrant cell lines (HepG2 and K562), the GM12878 cluster in a clade with two GM12878 bulk WGBS samples, and the Primary Fibro. cluster in a clade with two primary forearm fibroblast bulk WGBS samples (Fig. 2e, Supplementary Fig. 8,9). The confident assignment of these clusters for groups with as few as 10 cells (HEK293),suggests that sci-MET produces high quality profiles of DNA methylation, and paves the way for the assessment of tissues of complex cell type compositions.

**Figure 2.**
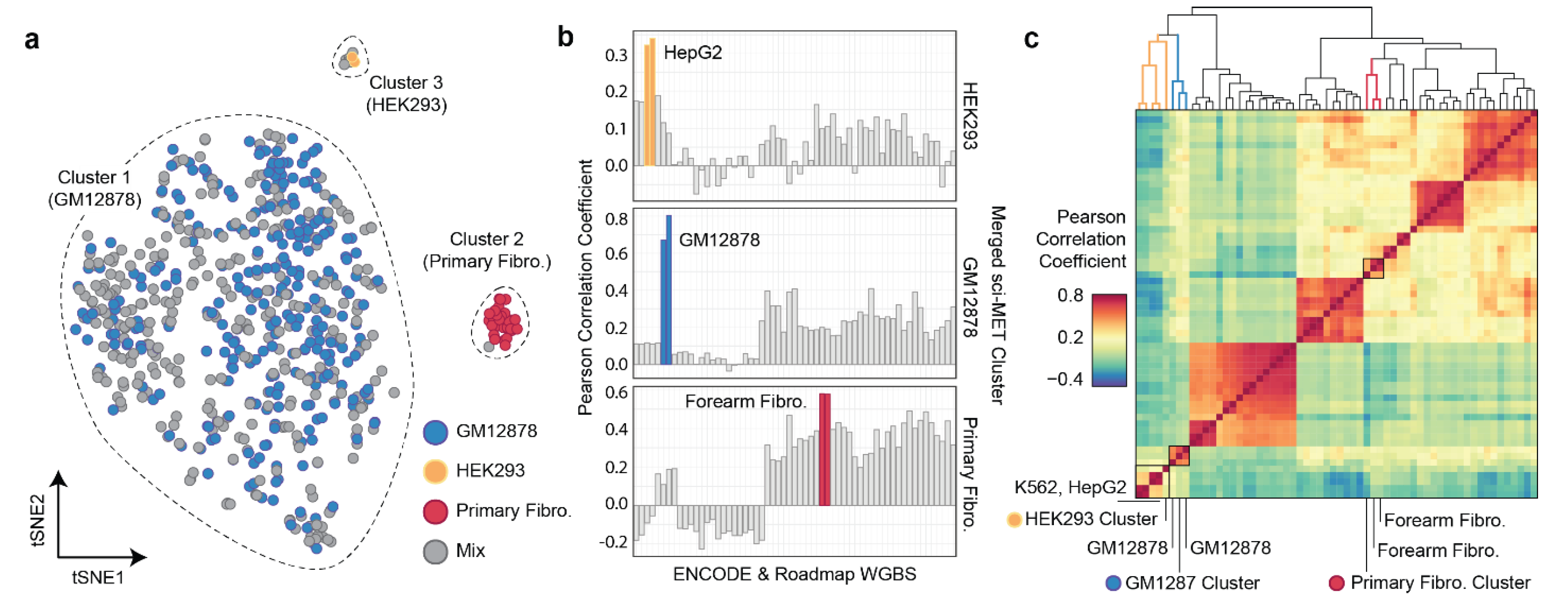
Sci-MET splits out single cell methylomes by cell type. **a.** Non-negative matrix factorization and t-distributed stochastic neighbor embedding (NMF-tSNE) was used to project single cell methylomes in twodimensional space (restricted to autosomal loci). Density based clustering was used to identify clusters. **b.**Single cell methylomes were aggregated over the three clusters and then correlated with publically available whole genome bisulfite sequencing data using the top 1,000 most variable regions in the Ensembl Regulatory Build. The two most highly correlated samples are in color. Note: For HEK293, no high coverage publically available WGBS data exists. **c.**Hierarchical clustering on the Pearson correlation values placed HEK293 in a clade with other cell lines (HepG2 and K562), GM12878 in a clade with WGBS of GM12878, and the Primary Fibro. cluster in a clade with two primary forearm fibroblast WGBS samples. Described clades are highlighted in color.

To our knowledge, sci-MET is the first molecular technique to produce single-cell whole genome bisulfite sequencing libraries at an order of magnitude scale improvement compared to current single-cell methods. Inherent in our protocol is the ability to scale to far greater numbers by expanding the number of indexes ***(e.g.*** 384 × 384; Supplementary Fig. 10). This puts sci-MET throughput on par with other high-throughput single cell assays. In addition to the increased throughput, we were able to achieve substantially improved read alignment rates when compared to existing low-throughput approaches, thereby dramatically reducing the sequencing burden for such studies. Our platform achieves both the throughput and cost-effectiveness that is required to scale single-cell DNA methylation assessment to levels comparable to other epigenetic and transcriptional properties and prompts the inclusion of this vital mark in large scale efforts to map all cell types in the body.

## Acknowledgements

We would like to thank Brooke DeRosa for culturing the primary fibroblast cell line for this project. We would like to thank other members of the Adey Lab for helpful suggestions and dialogue pertaining to this work, particularly Sarah Vitak. A.C.A. is supported by the Knight Cardiovascular Institute and the Medical Research Foundation of Oregon. B.J.O. is supported by a fellowship from the Sloan Foundation.

## Author Contributions

A.C.A. and R.M.M. conceived the sci-MET assay. R.M.M. carried out all sci-MET preparations with contributions from A.J.F. A.C.A., R.M.M., F.J.S., D.P., and S.N. designed the sci-MET adaptors and primers and reduced the assay to practice. R.M.M., F.J.S., D.P., and S.N. carried out all sequencing. R.M.M. led the data analysis. D.S. and Z.X. performed the NMF-tSNE analysis. K.A.T. provided additional analyses. F.J.S., J.S., C.T., and B.J.O. contributed to analysis design and edited the manuscript. A.C.A. supervised all aspects of the study. All authors approved the manuscript.

## Datasets Used

Publically available datasets used in this study were obtained from the ENCODE data portal with the following accessions: ENCFF039JFT, ENCFF092FNE, ENCFF103DNU, ENCFF110AZO, ENCFF121ZES, ENCFF122LEF, ENCFF157POM, ENCFF168HTX, ENCFF171ZRJ, ENCFF200MJQ, ENCFF210XTE, ENCFF215CKT, ENCFF216DJL, ENCFF241AQC, ENCFF247ILV, ENCFF256WDR, ENCFF257GGV, ENCFF186EKM, ENCFF517AOL, ENCFF545MIY, ENCFF266NGW, ENCFF279HCL, ENCFF315ZJB, ENCFF366UWF, ENCFF390OZB, ENCFF435SPL, ENCFF455TQO, ENCFF477AUC, ENCFF477GKI, ENCFF479QJK, ENCFF487XOB, ENCFF497YOO, ENCFF500DKA, ENCFF510EMT, ENCFF526PFA, ENCFF534RNT, ENCFF536RSX, ENCFF553HJV, ENCFF575GIN, ENCFF588IUK, ENCFF601NBW, ENCFF625GVK, ENCFF684JHX, ENCFF699GKH, ENCFF714SUO, ENCFF721JMB, ENCFF730NQT, ENCFF731IOY, ENCFF774GXJ, ENCFF795DNO, ENCFF831OYO, ENCFF835NTC, ENCFF837SXM, ENCFF847OWL, ENCFF867JRG, ENCFF874GGB, ENCFF913UZU, ENCFF918PML, ENCFF959WCA, ENCFF001SUN, ENCFF001SUL, ENCFF001SUG, ENCFF001SUD, ENCFF001SUJ, ENCFF001SUE, ENCFF001WYF, ENCFF001WYH, ENCFF001WYB, ENCFF001WYE, ENCFF001SUF, ENCFF001SUM, ENCFF001SUO, ENCFF001WYJ, ENCFF001WYK, ENCFF001SUI, ENCFF001SUP, ENCFF001SUQ, ENCFF741JQL, ENCFF549RWR, ENCFF323ZIV, ENCFF963GBQ, ENCFF800JNP, ENCFF001XDF, ENCFF639BKP, ENCFF363SIQ, ENCFF590RFP, ENCFF001WVZ, ENCFF001WWC, ENCFF001XHL, ENCFF001XHM, ENCFF001WWD, ENCFF001WWE, ENCFF001WWH, ENCFF001WWI, ENCFF523KSP, ENCFF066BAT, ENCFF825UAX, ENCFF765FCQ, ENCFF464QPC, ENCFF594VZB, ENCFF451UZW, ENCFF993MZN, ENCFF041SJL, ENCFF631QJF, ENCFF907IMB, ENCFF668WID, ENCFF050JWX, ENCFF418RFY, ENCFF483QXH, ENCFF301UTR, ENCFF019MRX, ENCFF715BRM, ENCFF367STH, ENCFF388TAT, ENCFF231GIV, ENCFF637XZK, ENCFF467BCP, ENCFF985WXP, ENCFF001VDK, ENCFF581RTT, ENCFF342JBJ, ENCFF037SXA, ENCFF422AIH, ENCFF001XCF, ENCFF001XCG, ENCFF127YXW, ENCFF498ERO, ENCFF781HLM, ENCFF046YRR.

## Methods

### Preparation of Unmethylated Control DNA

100 ng of unmethylated Lambda DNA (Promega, Cat. D1521) was treated with 4uL of 500 nM transposase-adaptor complex (transposome) pre-loaded with cytosine-depleted custom oligonucleotides in 10 uL of 1X Nextera Tagment DNA (TD) buffer from the Nextera DNA Sample Preparation Kit (Illumina, Cat. FC-121-1031) diluted with NIB to simulate reaction conditions for nuclei. Following incubation for 20 min at 55 °C, this reaction was cleaned with QIAquick PCR Purification Kit (Qiagen, Cat. 28104) and eluted in 30 uL of 10 mM Tris-Cl solution (pH 8.5). The tagmented, cleaned DNA was then quantified via Qubit 2.0 Flourometer dsDNA High Sensitivity Assay (Thermo Fisher, Cat. Q32854).

### Tissue Culture

Tissue culture cell lines (GM12878, Coriell; NIH/3T3, ATCC CRL-1658; HEK293, ATCC CRL-1554; Primary Fibro., inguinal fibroblast, GM05756, Coriell) were cultured in 5% CO2 at 37°C. GM12878 cells were grown in Roswell Park Memorial Institute media (RPMI, Gibco, Cat. 11875093) supplemented with 15% (by volume) fetal bovine serum (FBS, Gibco, Cat. 10082147), 1X L-glutamine (Gibco, Cat. 25030081), 1X Penicillin-Streptomycin (Gibco, Cat. 15140122), and gentamicin (Gibco, Cat. 15750060). HEK293 cells were grown in Dulbecco’s Modified Eagle’s media (DMEM, Gibco, Cat. 11995065), supplemented with 10% FBS, and 1X L-glutamine. NIH/3T3 cells were grown in the same preparation of DMEM as HEK293 cells. Primary Fibroblasts were cultured in a growth medium comprised of DMEM/F12 (with GlutaMax; Thermo Fisher), 10% fetal bovine serum (FBS; Thermo Fisher), 1% MEM Non-Essential Amino Acids (Thermo Fisher), and 1% Penicillin/Streptomycin (Thermo Fisher). Adherent cell lines were grown to ~90% confluency at the time of harvest.

### Sample preparation and nuclei isolation

For library preparation, cells were pelleted if cultured in suspension, or trypsinzed (Gibco, Cat. 25200056), if adherent. Cell were washed once with ice cold PBS and carried through cross-linking (for the xSDS method) or directly into nuclei preparation using nuclei isolation buffer (NIB, 10mM TrisHCl pH 7.4, 10 mM NaCl, 3mM MgCl2, 0.1% Igepal, 1X protease inhibitors (Roche, Cat. 1187358001)).

### Nucleosome Depletion

Detailed step-by-step protocol for nucleosome depletion and all subsequent steps can be found in the Supplementary Protocol. Nucleosome depletion and combinatorial indexing strategies were performed similar to previously described, with some variations^20^.

*Lithium-assisted nucleosome depletion (LAND)* was performed for generation of GM12878-only and Human/Mouse libraries. Prepared nuclei were pelleted and resuspended in NIB supplemented with 200 uL of 12.5 mM lithium 3,5-diiodosalicylic acid (Sigma, Cat D3635) for 5 minutes on ice before addition of 800 uL NIB and then taken directly into the combinatorial indexing protocol.

*Cross-linking and SDS nucleosome depletion (xSDS).* Cells were cross-linked by incubation in 10 mL of media with 1.5% formaldehyde (final conc. by vol.) and incubated at room temperature for 10 minutes with gentle agitation. Cross-linking was quenched with 800 uL 2.5 M glycine and incubated on ice for 5 minutes. Cells were then spun down, washed with ice-cold PBS, and resuspended in ice cold NIB for a 20 minute incubation on ice with gentle agitation. Cells were then pelleted, washed with 900 uL of 1X NEBuffer 2.1m and resuspended in 800 uL 1X NEBuffer 2.1 with 0.3% SDS (Sigma, Cat. L3771) and incubated at 42 °C with vigorous shaking for 30 minutes in a thermomixer (Eppendorf). 200 uL of 10% Triton-X was added to quench, and the solution was incubated at for another 30 minutes at 42 °C with vigorous shaking. Nuclei were then taken into the combinatorial indexing protocol. We were concerned that the crosslinking may affect the bisulfite conversion reaction; however, based on the methylation rates (particularly for those of nonCG methylation which were very low in concordance with expectations), we determined that not to be the case.

### Combinatorial indexing via tagmentation

Nuclei were stained with 8uL of 5mg/mL DAPI (Thermo Fisher, Cat. D1306) and passed through a 35-um cell strainer. A 96-well plate was prepared with 10 uL of 1X TD buffer diluted with NIB in each well. Fluorescence-assisted nuclei sorting (FANS) was performed with a Sony SH800 flow sorter to sort 2,500 single nuclei into each well in fast sort mode (Supplementary Fig. 10). 4uL of 500 nM transposome, pre-loaded with cytosine-depleted, uniquely indexed, custom oligonucleotides was placed in each well (described in supplement, transposomes assembled as described in Amini ***et. al.*** 2014, ref.^17^). Reactions were incubated at 55°C for 20 minutes. All wells were then pooled and stained with DAPI as done for the first FANS sort. A second 96-well plate was prepared with each well containing digestion reagents as described by the manufacturer’s protocol for the EZ-96 DNA Methylation MagPrep Kit (Zymo, Cat. D5040) at one-fifth the volumes (for 5 uL per well). 22 post-tagmentation nuclei from the pool of all reactions were sorted into each well using the single-cell sorting setting. Some wells were selected to receive only 10 nuclei, to allow for unmethylated controls. The plate was then spun down at 600 x g for 5 minutes at 4°C.

### Library preparation

Prior to bisulfite conversion, several wells, which only received 10 nuclei in the final sort, were spiked with ~35 pg of the prepared unmethylated control DNA, to keep DNA mass constant per well. Nuclei were then processed following manufacturer’s protocol for the EZ-96 DNA Methylation MagPrep Kit, with volumes reduced to one-fifth those described by the manufacturer to allow for single-well reaction processing, and other slight modifications. Following the final post-bisulfite library cleanup, each well was eluted in 25 uL of Zymo M-Elution Buffer and transferred to a well in a 96-well plate prepared with the following reaction mixture for linear amplification: 16 uL PCR-clean ddH2O, 5 uL 10X NEBuffer 2.1 (NEB, Cat. B7202), 2 uL 10 mM dNTP mix (NEB, Cat. N0447), and 2 uL of 10 uM random nonamer primer with a partial sequence of the Illumina Standard Read 2 sequencing primer (9NP, 3’- NNNNNNNNNAGATCGGAAGAGCACACGTCTG-5’). To render libraries single stranded prior to linear amplification, reactions were heat shocked at 95°C for 45 seconds and then flash cooled on ice. Following cooling, 10 U Klenow (3’->5’ exo-) polymerase (Enzymatics, Cat. P7010-LC-L), was added to each reaction and followed by incubation at 4 °C for 5 minutes, then a slow ramp of +1 °C/15 seconds, and 37 °C for 90 minutes. This was repeated for 2-4 times dependent on library and in accordance with previously describe scWGBS protocols (Supplementary Fig. 1)^12^. For each repetition, 1 uL 10 uM 9NP, 1 uL 10 mM dNTP mix, 1.25 4X NEBuffer 2.1, and 10 U Klenow (3’ -> 5’ exo-) polymerase was added after the heat shock and cooling. Following completion of linear amplifications, wells were cleaned with 1.1X (by volume) of 18% PEG SPRI Bead mixture (Sera-Mag SpeedBeads (GE, Cat. 65152105050250) washed and resuspended in 18% PEG 8000 (by mass), 1M NaCl, 10mMTris-HCl, pH 8.0, 1mM EDTA, 0.05% Tween-20), with a 5 minute room temperature incubation, then placed on a magnetic rack until the supernatant was cleared. The supernatant was discarded, and beads were washed with 80% ethanol while held in place by the magnets. Beads were then dried and libraries were eluted in 21 uL 10 mM Tris-Cl (pH 8.5). The full 21 uL eluate was then placed into a 96-well plate prepared with a PCR reaction mixture containing 25 uL 2X KAPA HiFi HotStart ReadyMix (Kapa, Cat. KK2602),2 uL of 10 uM forward and reverse uniquely indexed primers (each introducing a 10-nt indexing sequence), and 0.5 uL of 100X SYBR Green I (FMC BioProducts, Cat. FC-121-1031). Real time PCR was performed on a Bio-Rad CFX thermocycler with the following conditions: 95°C for 2 minutes, (94°C for 80 seconds, 65°C for 30 seconds, 72°C for 30 seconds [Image]) for 18-22 cycles. PCR was stopped once libraries reached the inflection point of measured SYBR green fluorescence. Following PCR, libraries were then pooled by column (10 uL/well) and with 0.8X (by volume) 18% PEG SPRI Bead Mixture as described previously. Libraries were eluted off the magnetic beads in 25 uL of 10 mM Tris-Cl (pH 8.5).

### Library quantification and sequencing

Libraries were pooled and quantified between the range of 200 bp and 1 kbp using a 2100 Bioanalyzer DNA High Sensitivity kit (Agilent, Cat. 5067-4626; Supplementary Fig. 11). Pools were sequenced on either an Illumina NextSeq 500, HiSeq 2500 or HiSeq X, loaded at 0.9 pM, with a 5% PhiX spike-in to improve complexity (or 30% spike-in for the NextSeq 500). All sequencing runs used a custom locked-nucleic acid (LNA) oligonucleotides for custom sequencing primers to match the standard chemistry temperatures (Supplemental Fig. 1). With the exception of the first GM12878-only library pool, libraries were sequenced with a custom sequencing chemistry protocol (Read 1: 100 imaged cycles; Index Read 1: 10 imaged cycles, 27 dark cycles, 11 imaged cycles; Index Read 2: 10 imaged cycles).

### Sequence read processing

Reads were processed using bcl2fastq (Illumina Inc., v2.19.0) with the “--create–fastq–for–index–reads” and “--with–failed–reads” options to produce fastq files. Fastq reads were then identified by indexes, requiring each index (the two 10-nt indexes introduced by PCR, and the 11-nt index introduced by tagmentation) to independently be within a Hamming distance of two from the expected reference sequences. Reads with all three indexes assigned had the respective reference index sequences concatenated to a barcode and appended to the read name, which served as the cell identifier. Reads were then trimmed using TrimGalore! (v0.4.0) with option “-a AGATCGGAAGAGC” to identify adapters. Trimmed reads were quality checked using FastQC (v0.11.3) for adapter content, percent base across reads for bisulfite conversion biases, and k-mer bias. Alignment to the human (GRCh37), mouse (GRCm38), or a combined human-mouse hybrid genome was performed with Bismark (v0.14.3) using “--bowtie2” and “--unmapped” options^25^. Aligned reads were then de-duplicated based on barcode, chromosome, and starting position.

### GM12878-only library development

GM12878-only libraries were generated as described above with alterations/specifications as follows: library were generated using the LAND method for nucleosome depletion, libraries were generated using four rounds of linear amplification, and were sequenced in a paired-end manner. For the paired-end sequencing strategy the following custom sequencing chemistry protocol was used (Read 1: 50 imaged cycles; Index Read 1: 10 imaged cycles, 27 dark cycles, 11 imaged cycles; Index Read 2: 10 imaged cycles; Read 2: 50 imaged cycles). Sequencing reads were processed using slightly modified read processing pipeline. Trimming was performed with TrimGalore! using the “-paired” option, we observed biases at the start of both read 1 and read 2 sequences, likely due to the random priming strategy, and consequently trimmed the reads with options “--clip_R1 6”, “--clip_R2 9”. We aligned reads to the GRCh37 reference genome with Bismark with an added “-p” option for the paired-end alignment.

### Human-mouse library development

Human (GM12878) and mouse (NIH/3T3) cell lines were mixed following nuclei isolation, but before nucleosome depletion in a roughly equal ratio. Nucleosomes were then depleted using the LAND technique and processed as described above. Reads were aligned to a hybrid human-mouse genome to estimate barcode collision rate. We observed an increase in reads mapping to either human or mouse chromosomes dependent on read-depth, likely reflecting the lack of specificity in alignment of bisulfite sequencing data. To estimate barcode collision rate we identified putative single cell libraries with < 90% of reads that aligned to a single species which represents approximately half of the total collision rate. We also generated a second human-mouse library using a mixture of human (HEK293) and mouse (NIH/3T3) cells which underwent xSDS nucleosome depletion. The human-mouse xSDS library was processed as described above.

### Cell line deconvolution library development

To assess the ability of sci-MET to separate out different cell types using a low-coverage, high-cell count approach, we generated a library pool consisting of GM12878 (a B-lymphoblastoid cell line, 40%), HEK293 (a kidney epithelial cell line, 20%), and GM05756 (primary inguinal fibroblast line, 40%). Cell lines were brought through the sci-MET protocol via xSDS, both in parallel, and as an equal ratio mix combined after nuclei isolation. We suspect that this ratio was dramatically altered due to the FANS gating that we performed which likely excluded the majority of the aneuploid HEK293 cells which are difficult to distinguish from euploid doublets. Furthermore, for the majority of wells in which the cell identity was known, the cells were GM129878, thus likely favoring the FANS gating to that cell’s profile. It is important to note that this challenge would persist for any method of single cell profiling that requires single cell sorting, such as all of the existing single cell methylation assay platforms. Libraries were processed as described above.

### Single-cell Discrimination

We calculated the read threshold for including barcodes as individual cells by first performing k-means clustering (k=3) based on the log10 number of unique aligned reads per cell and then fitting a normal distribution to the cluster containing the cells with the highest number of unique aligned reads. The threshold was then defined based on the 95% confidence interval (CI) of the fitted normal distribution (mean-(1.96 x SD)). We used the kmeans function in R for clustering and the ***MASS*** (v. 7.3-45) and ***mixtools*** (v. 1.1.0) packages for fitting the normal distributions. Peaks did not show clear separation for the GM12878-only prep due to low coverage. As an alternative approach, ***mixtools*** was used to fit mixed normal distributions to the clustered data from which we calculated the 95% CI on the peak with the highest number of unique reads. (Supplementary Fig. 12).

### Quality Control

We assessed bisulfite conversion efficiency in our preparations through spike in of unmethylated lambda DNA. We aligned fastq reads with the respective 11-nt tagmentation index to the lambda genome (GenBank: J02459.1) using Bismark. We de-duplicated reads, and filtered to high quality alignments (≥Q30). We observed a highly efficient bisulfite conversion across sci-MET library constructions (>99%).

Individual barcodes per library were assessed for mapping efficiency (calculated as aligned reads/fastq reads assigned to a barcode), and complexity (calculated as de-duplicated, aligned reads/aligned reads assigned to a barcode). Our protocol for library construction both increased the throughput of single-cell generation, and significantly increased mapping efficiency compared to previously methods. We required cells which met read threshold cutoff to have a mapping efficiency of ≥%, a nonCG methylation of ≤ 5% for downstream clustering analysis. Of note, the theoretical maximum number of QC-passing sci-MET single cell libraries for an experiment is the number of transposase-indexed nuclei per bisulfite + PCR well multiplied by the total number of bisulfite + PCR wells. In the case of our human cell type mix experiment this results in a theoretical maximum of 844 libraries. 641 single cells of the possible 844 sorted passed read threshold and quality filters, yielding a success rate of 75.95%. We further stratified our library pool to assess the effect of various rounds of linear amplification on single-cell library quality. We found that four rounds of linear amplification significantly increased mapping efficiency (p-value = 7.83 x10^-16^, t= 8.27, Student's t test; Supplementary Fig. 2). Further we found that increased rounds of linear amplification increased the library complexity. We fitted two-factor saturation curves to single-cell libraries using the ***drc*** (v3.0-1) package's ***drm*** function in R dependent on rounds of linear amplification. For four rounds and three rounds of linear amplification, our projected upper asymptotes (full sequencing saturation) were 3.88 x 10^6^, and 2.71 x 10^8^ unique CGs per single cell library, respectively (Figure 1e).

### Coverage Bias across Annotations

We calculated the coverage bias in individual cells across DHS, CG Islands, and Histone (H2AFZ, H3K27ac, H3K36me3, H3K4me1, H3K4me3, H3K4me3,H3K79me2, H3K9ac, H3K9me3, H4K20me1, H3K27me3) sites using annotated DNase, methylation profiling and CHIP-seq peak data from the publically available UCSC and ENCODE databases^23,24^. We used bedtools multicov (v. 2.22.0) to determine the coverage for each cell across all sites of each annotation bed file. We then determined the fraction of total reads per kilobase pair (kbp) by summing the coverage across all sites in a cell and normalizing by the reads per cell and by the sum of the genomic distance of the peak sites (Supplementary Fig. 4).

### CG Sites Covered Per N Cells Analyzed

We simulated the number of unique CG sites covered in an experiment by an arbitrary number of cells using sciMET (Human cell types data) by performing 100 iterations of sampling of n=(10, 20, 30, 40, 50, 60, 70, 80, 90, 100, 150, 200,250, 300, 350, 400, 450, 500, 550, 600, 650) cells. We then calculated the aggregate number of unique CG sites covered across all cells for each sampling and fitted a lowess curve (using R package ***ggplot*** v.2.2.1) to the unique CG sites per n cells sampled saturation plot (Supplementary Fig. 5).

### Non-negative Matrix Factorization, tSNE, and clustering

Non-negative Matrix Factorization (NMF) is a non-supervised data decomposition technique^26^. Here we used NMF to learn new feature representations. NMF is mathematically approximated by: **A^*mxn*^**≈**W**^*mxk*^•**H**^*k×n*^, where **A** is the matrix representing the single cell methylation profiles of n samples. **W** is a dictionary matrix with a much smaller k<<m.**H** is the activation coefficients on the new basis. All the three of them are non-negative. The column vectors in **W** are called ***meta-feature,*** which are higher-level abstraction of the original methylation levels and each column in **H** is meta-expression on the new basis of each sample. Here we set ***k =*** 12 to get matrix **A**factorized into low-rank matrix **W** and **H**. In this way, we extracted the uncorrelated basis and the coefficient matrix H of the new basis by significantly reducing the dimension of the features. Since relatively few basis vectors are used to represent many data vectors (k<<m), good approximation can only be achieved if the basis vectors discover structure that is latent in the data, which will help the next sample clustering and visualization. Then, given the learned feature representation, Student t-Distributed Stochastic Neighbor Embedding (t-SNE) package ***Rtsne*** (v.0.13) for R is used to plot the meta-expression matrix **H**^*k×n*^ with default parameters. Clustering on the NMF-tSNE coordinates was performed using the Density Based Clustering of Applications with Noise (DBSCAN; v.1.1-1) with an epsilon value of 4 and a minimal cell seed threshold of 4, ref.^27^. This process was performed for cells with ≥ 30,000 unique aligned reads (Fig. 2c) as well as for just cells with ≥ 50,000 unique aligned reads (Supplementary Fig. 13), which provided no qualitative difference.

### Methylation over Genomic Annotations

Methylation rate over ChIP-seq and other genomic annotations was carried out by aggregating the methylation fractions in percentile windows for 5,000 bp upstream of the feature, through the feature set, and 5,000 bp downstream of the feature and smoothed over three percentile windows. Methylation rates were carried out for each individual cell as well as for the combination of cells of each specific sample type in the case of the human cell type mix experiment.

### Window Summaries and Correlations over Ensembl Regulatory Regions

We quantified methylation rate across Ensembl Regulatory Build windows using a previously described method^12^. Density plots of methylation rates per cell, and per reads collapsed by cluster identity were generated with ***ggplot2,*** demonstrating a strong expected bimodality even within low-coverage libraries. Using ENCODE and Epigenome Roadmap bulk wGBS samples, we quantified a weighted methylation rate and variance across samples using the Ensembl Regulatory Build loci^22^. We next took the top 1000 most variable loci across the bulk samples and summarized methylation rates within single-cell clusters identified above. We performed a Pearson correlation of methylation rates with the bulk WGBS samples using base R ***cor*** function. Biclustering was performed using the R package ***gplots*** (v. 3.0.1) ***heatmap2*** function.

---

**Supplementary Figure 1.**
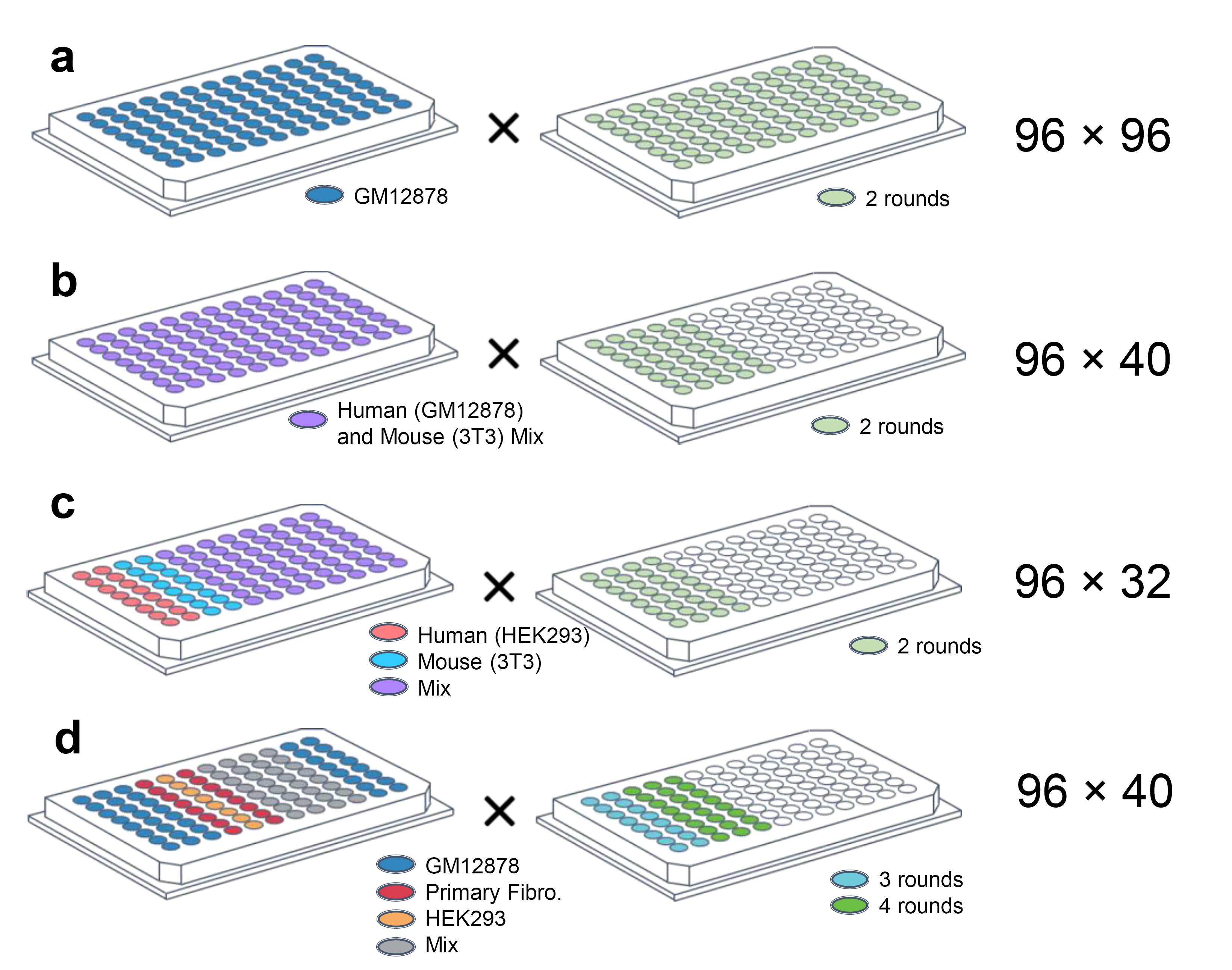
Summary of experiments and indexing design. **a.** GM12878 initial experiment using lithium-based nucleosome depletion. **b.** Human and mouse mix experiment using lithium-based nucleosome depletion. **c.** Human and mouse mix experiment using crosslinking and SDS nucleosome depletion. **d.** Human cell type mixing experiment using crosslinking and SDS nucleosome depletion. Rounds indicates the number of rounds of linear amplification post-bisulfite treatment. The 96 × N values indicate the scale of the experiment as described in the main text.

**Supplementary Figure 2.**
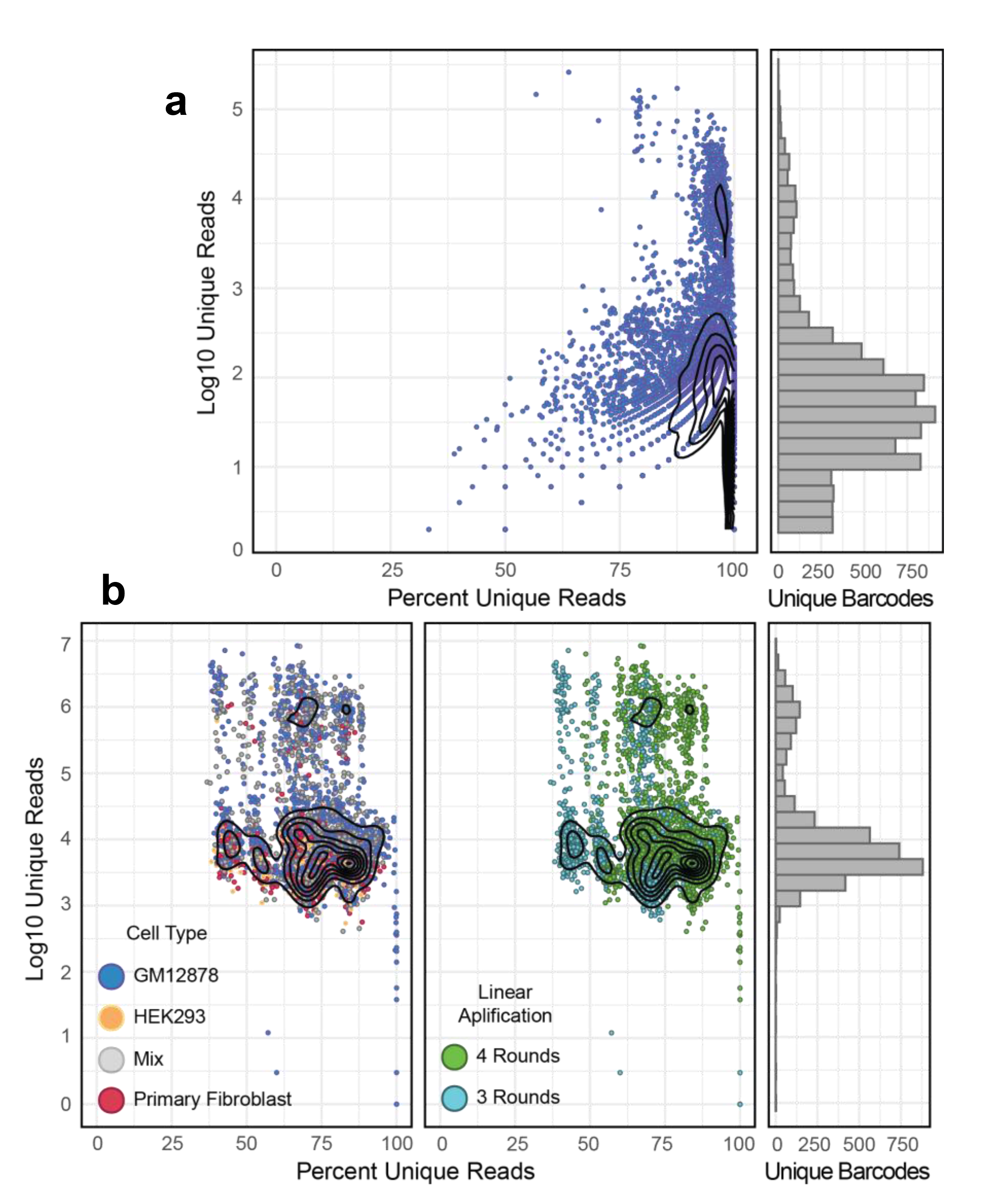
Library read count and unique read percentages for (**a**) the initial GM12878 only experiment which was sequenced to low depth, and (**b**) the human cell line mix experiment. X-axis indicates the percent of unique reads out of total aligned reads, Y-axis is the log_10_ unique read count for each cell (each represented as a point). The high read count distribution indicates true single cell libraries and the lower distribution is the background noise. In the middle panel of **b**, rounds indicates the number of rounds of linear amplification.

**Supplementary Figure 3.**
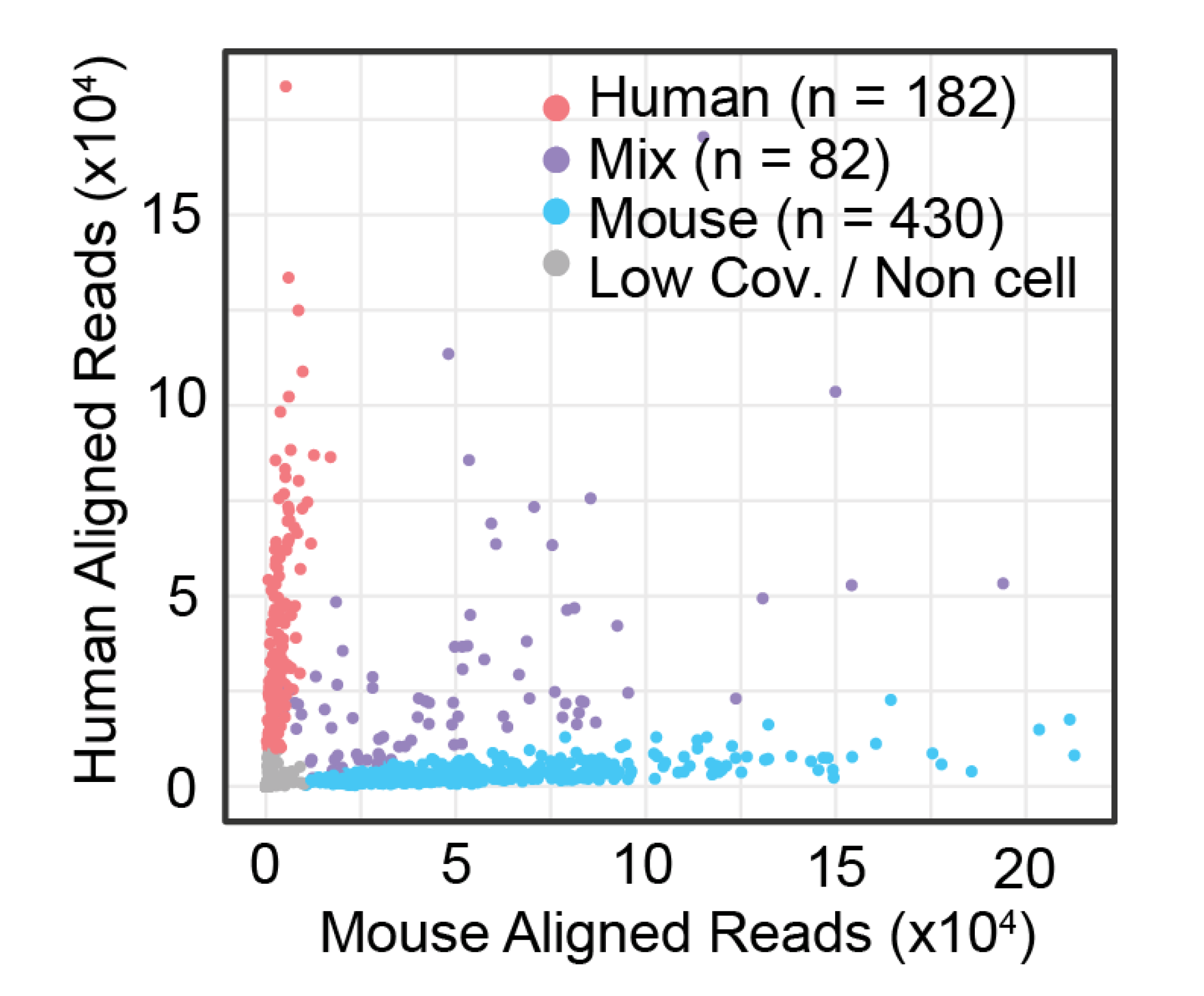
Sci-MET assay on a mix of human and mouse cells using lithium-based nucleosome depletion. A high collision rate (22% total estimated collision rate) was observed, possibly due to library molecule leakage after transposition.

**Supplementary Figure 4.**
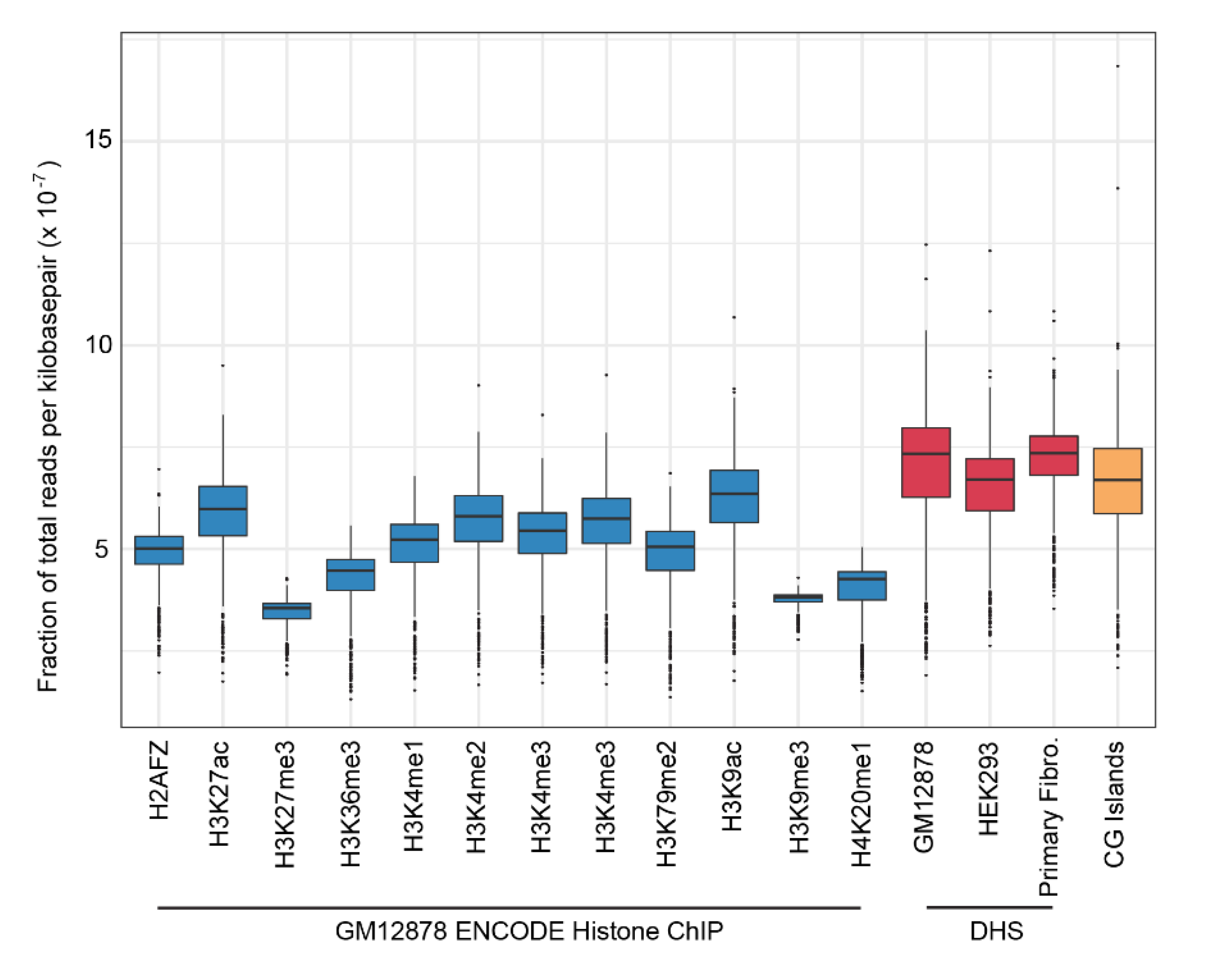
Bias of coverage for sci-MET libraries constructed for the cell type mix experiment. Each cell had its read count within feature windows normalized to the total count for the cell and the total combined size of the features. DHS= DNase-seq hypersensitivity. All ChIP-seq and DHS feature sets were obtained from the ENCODE data portal.

**Supplementary Figure 5.**
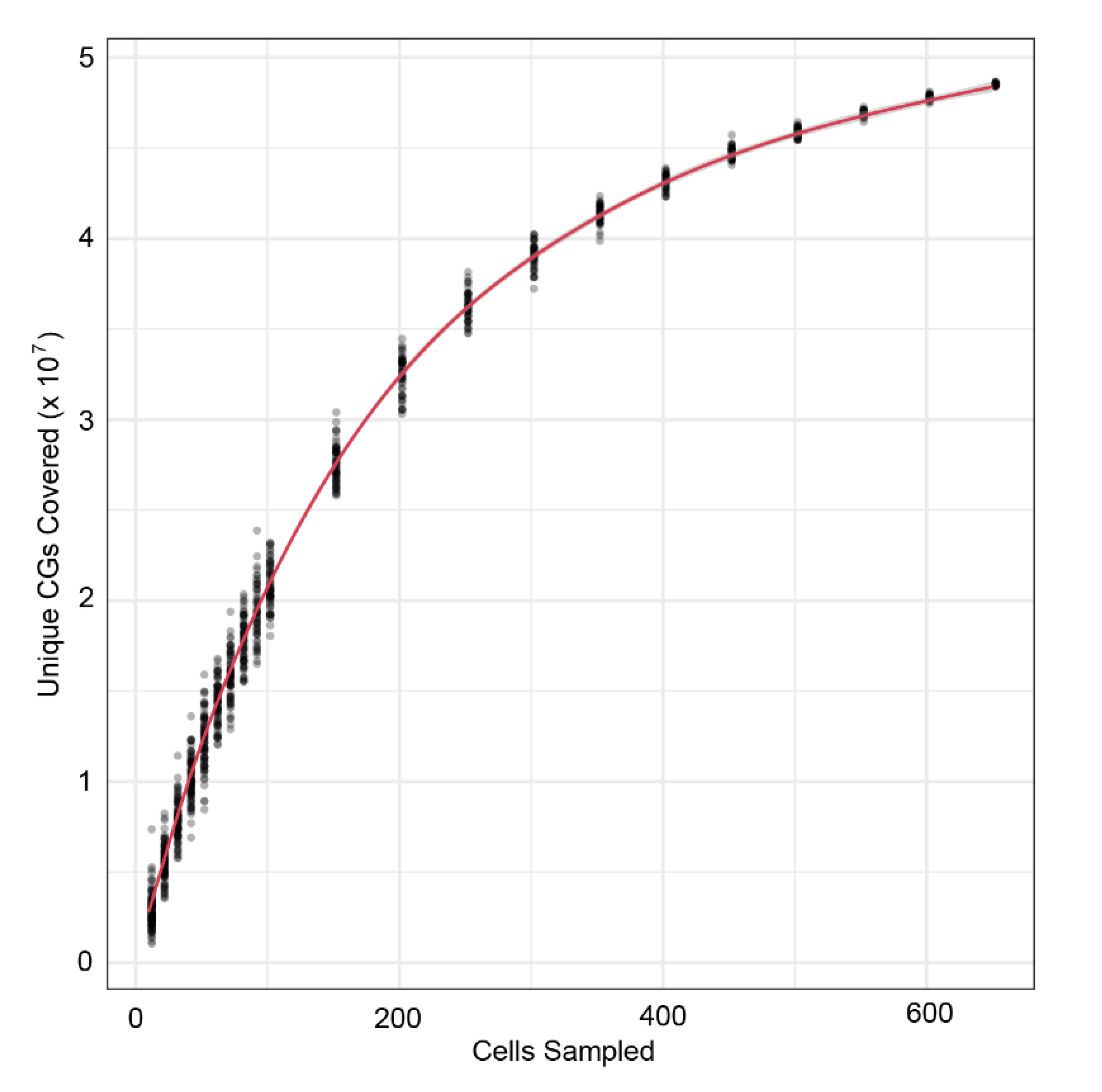
Total unique CG dinucleotides covered as a function of the number of sciMET single cell libraries merged. 100 random subsets of cells were sampled at each increment from the human cell type mix experiment for cell libraries containing a minimum of 30,000 unique aligned reads.

**Supplementary Figure 6.**
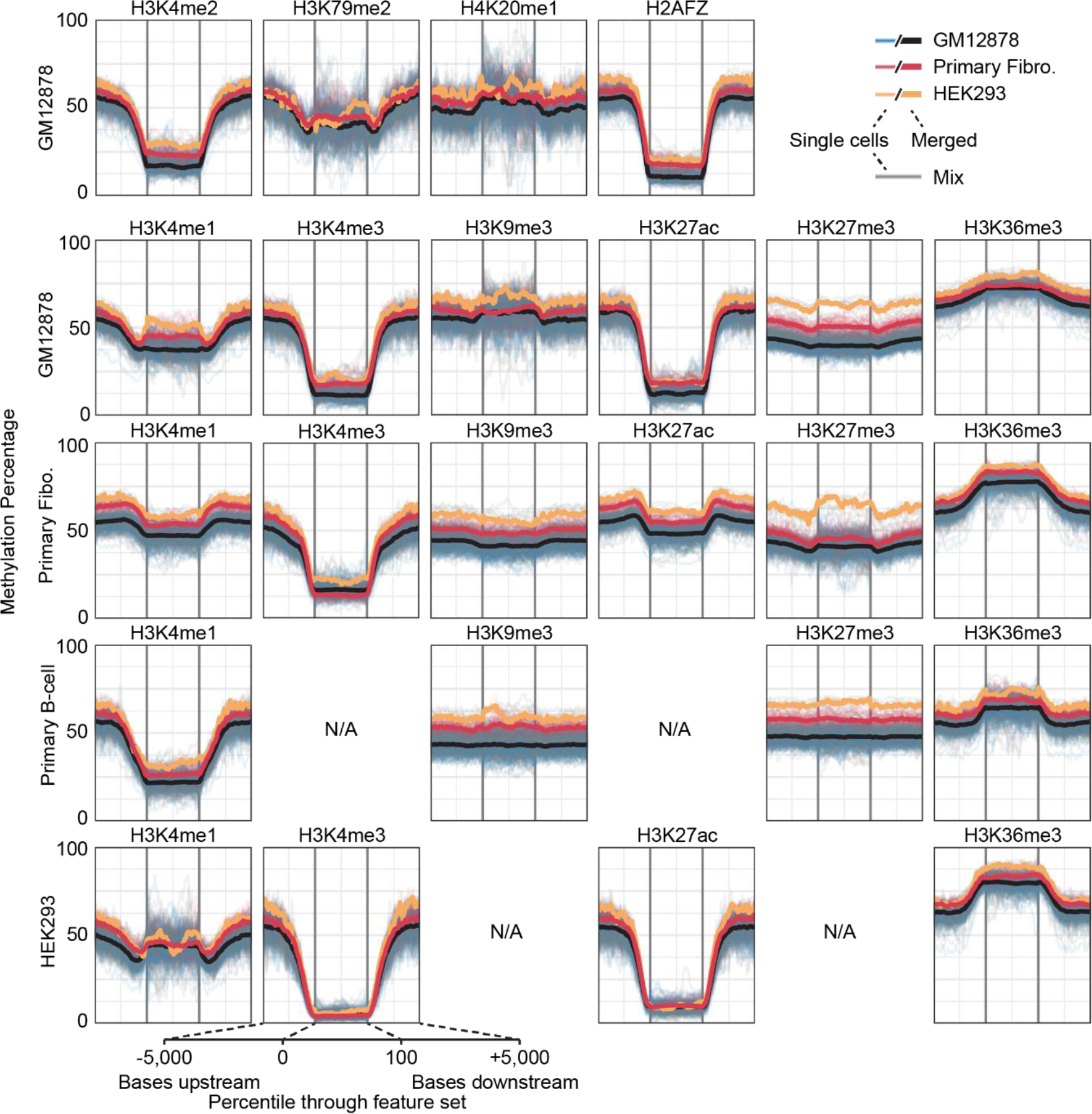
Methylation rate aggregated over ENCODE features identified by ChIP-seq. Methylation rate was calculated for the upstream 5,000 bp of each feature, throughout the feature (defined as percentile of progression through the feature), and 5,000 bp downstream and then aggregated across all features for each single cell (transparent lines) and then averaged across the pure cell types (solid lines).

**Supplementary Figure 7.**
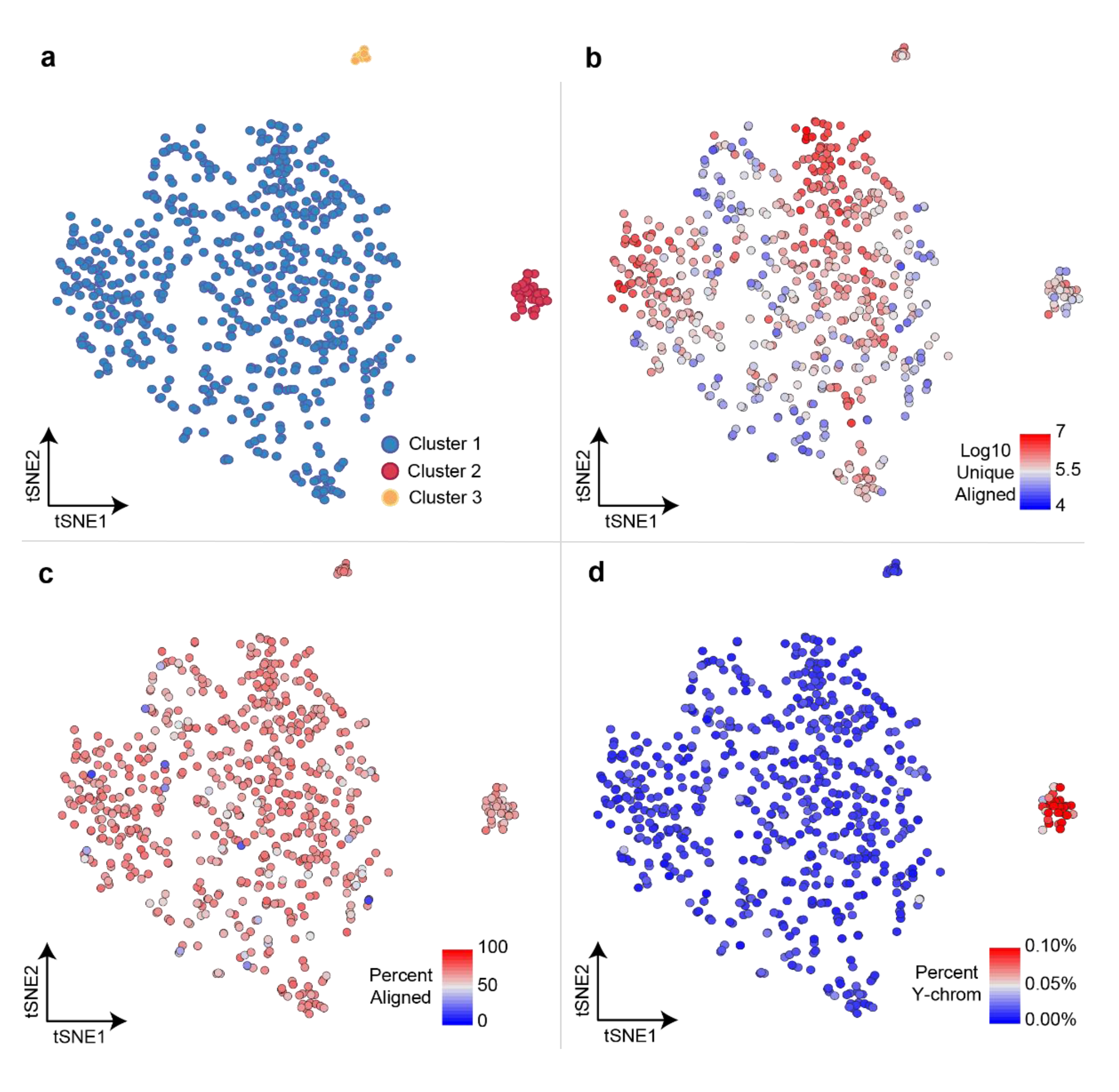
Clustering and library metrics. **a.** After NMF-tSNE, cell coordinates were used to identify clusters using the density-based clustering algorithm dbscan with an epsilon value of 4 and a minimum of 4 cells per seed. Clusters are colored based on their corresponding cell types. **b.** Log10 unique aligned reads with an alignment quality score ≥ 10. **c.** Percent of reads aligned for each library. **d.** Percent of reads that align to the Y-chromosome. BJ cells (foreskin fibroblast cell line) are the only cells derived from a male and form the far right cluster.

**Supplementary Figure 8.**
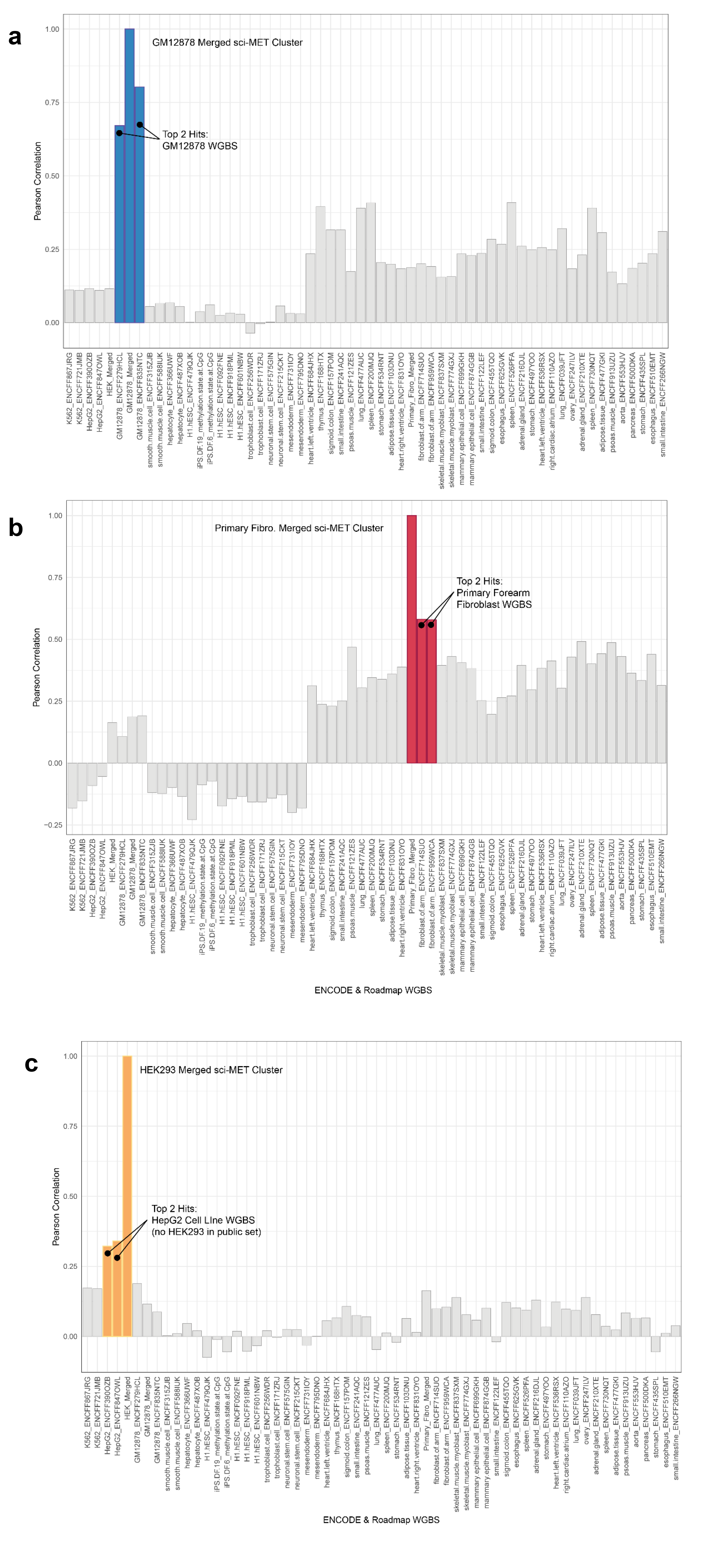
Correlation of aggregate clusters to ENCODE WGBS data sets for the top 1,00 most variable Ensembl regulatory regions for the (a) GM12878, (b) Primary Fibro., and (c) HEK293 merged cell clusters. Datasets are in the order of the hierarchical clustering.

**Supplementary Figure 9.**
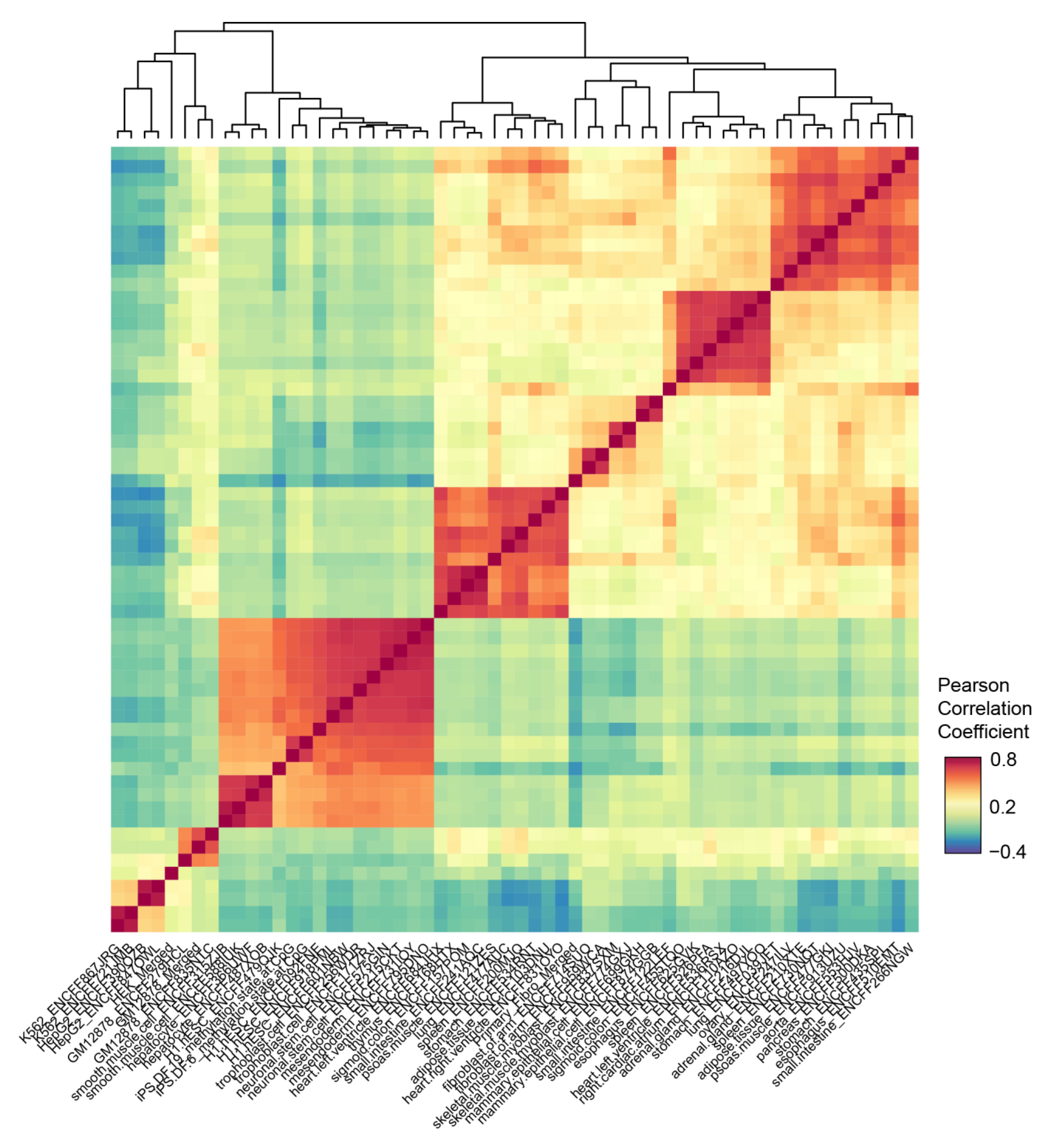
Biclustering of correlation coefficients for publically available WGBS data and the merged sci-MET clusters of the top 1,000 most variable regions in the Ensembl regulatory build. Of note, the GM12878 sci-MET cluster is in a clade with two GM12878 WGBS samples, the HEK293 sci-MET cluster is in a clade with HepG2 and K562 cell lines (there is no HEK293 public dataset used), and the Primary Fibroblast sci-MET cluster is in a clade with two primary forearm fibroblast WGBS samples.

**Supplementary Figure 10.**
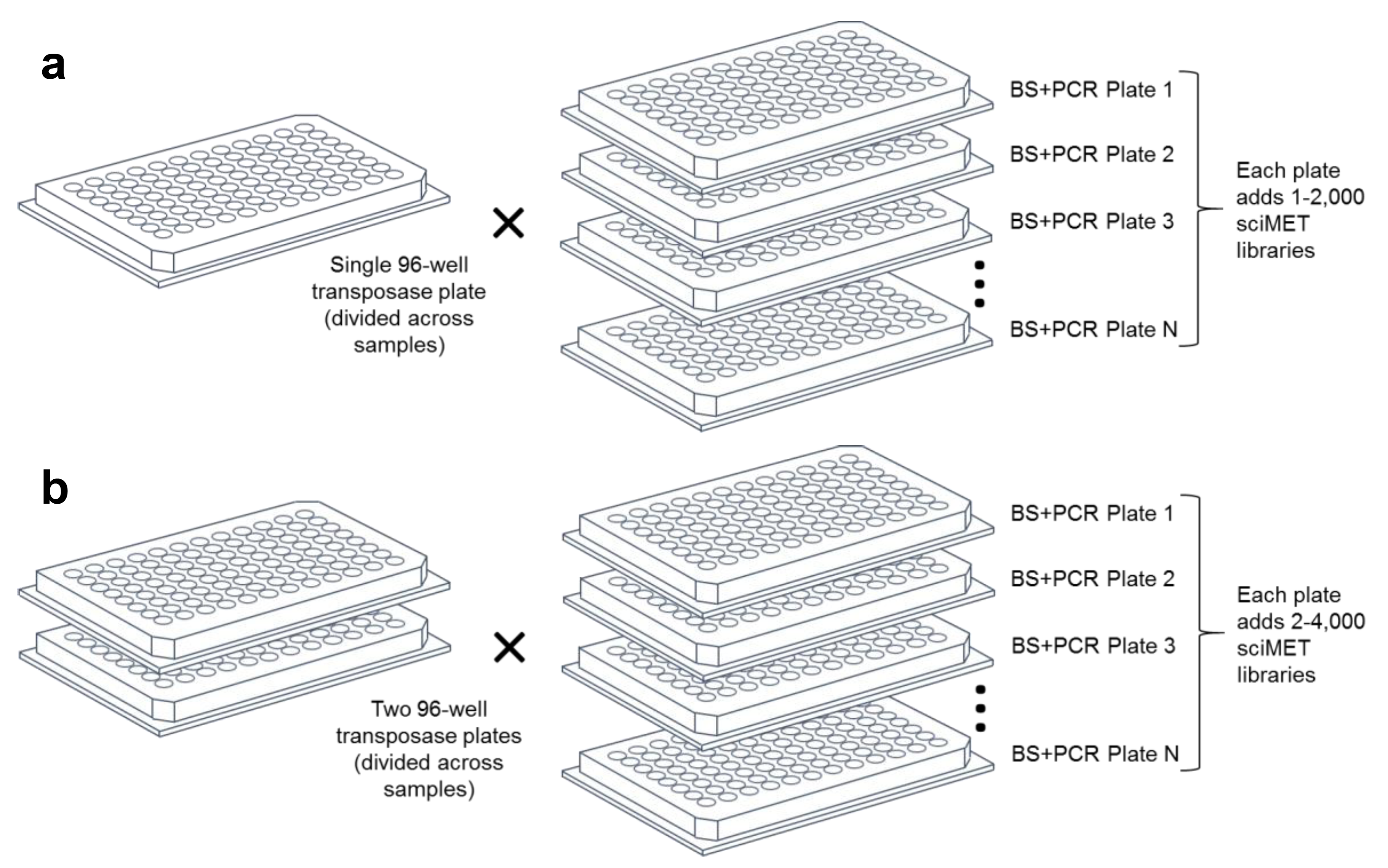
Scalability of sciMET platform. a. Scalability of sciMET using a single initial 96-well plate of indexed transposomes. b. Scalability of sciMET when increasing the number of initial indexes which increases libraries produced for each subsequent plate. The expected cell count can be represented as:

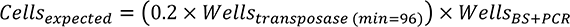

**Supplementary Figure 11.**
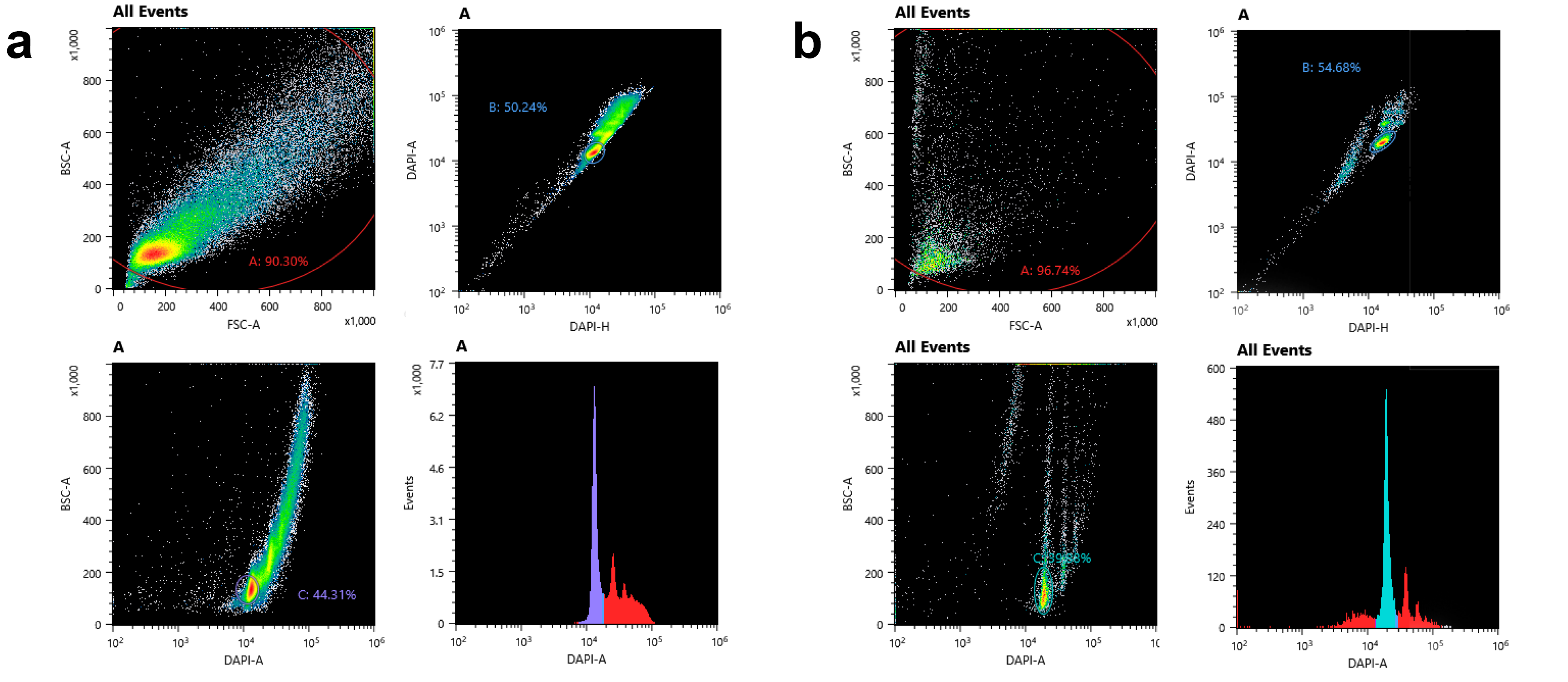
FANS sorting for a sci-MET prep. **a.** Initial sort of nuclei into wells of the transposase plate. **b.** Second sort of tagmented nuclei into the bisulfite reaction and subsequent PCR.

**Supplementary Figure 12.**
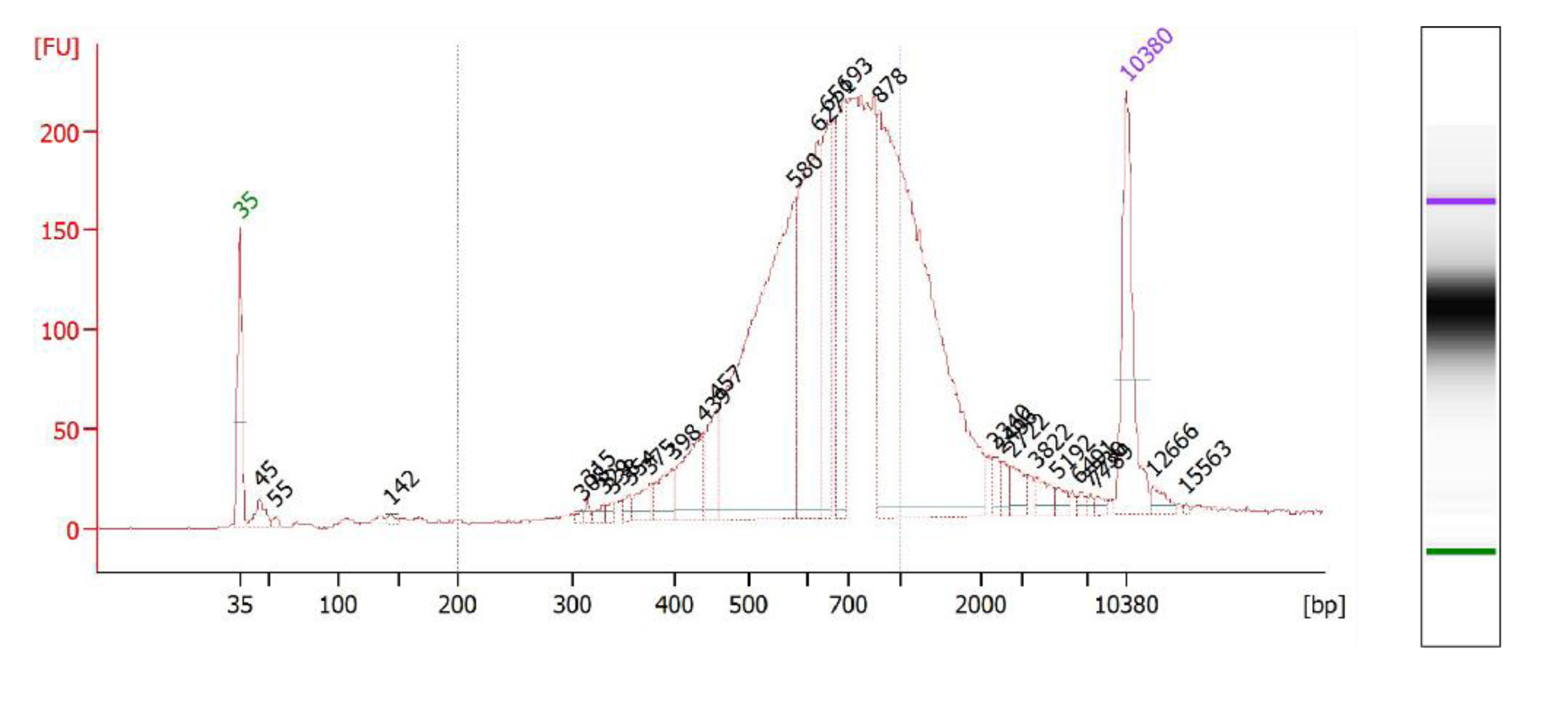
Bioanalyzer trace of the human cell line mix library.

**Supplementary Figure 13.**
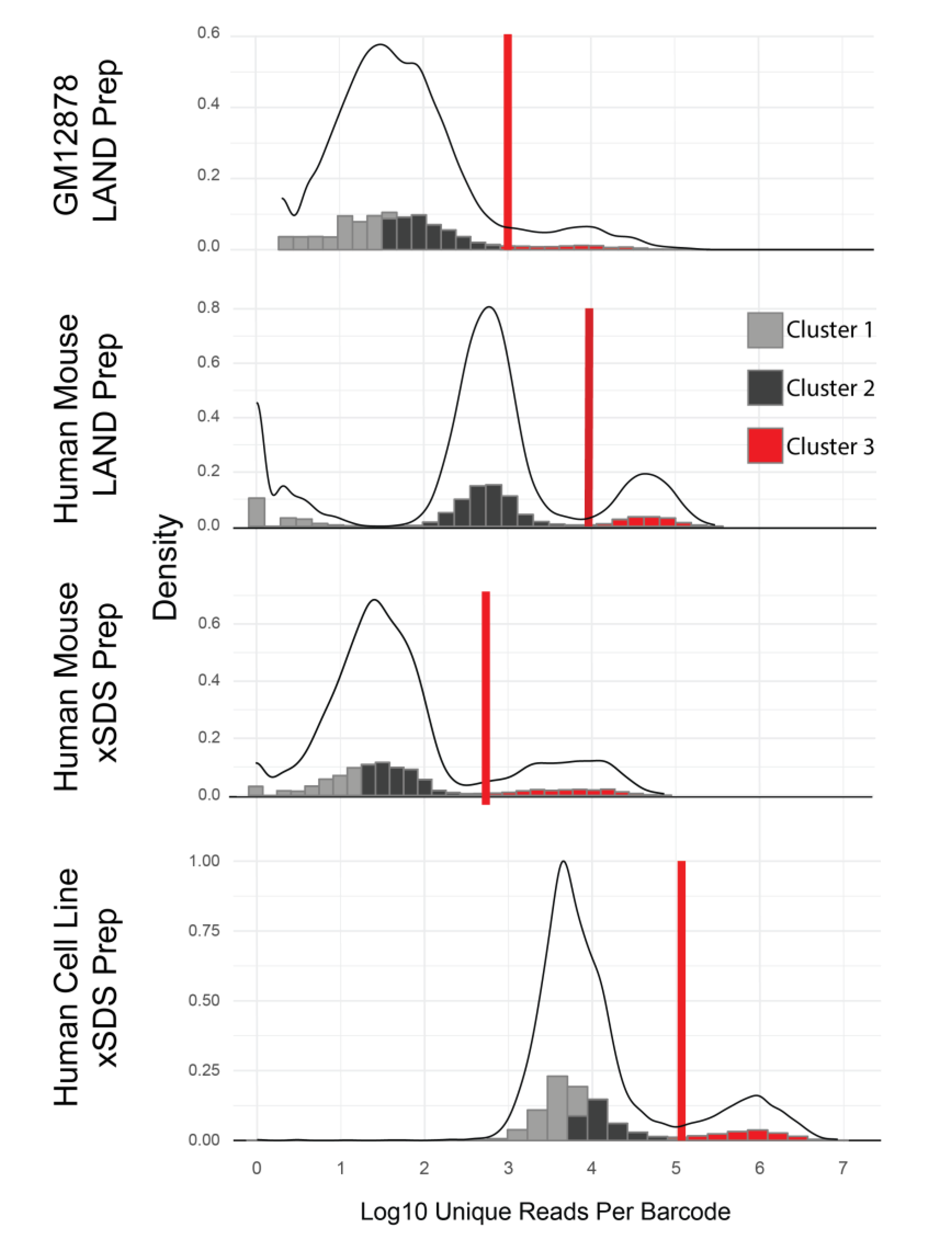
Discrimination of single cell libraries through library read counts. Histograms and density plots of the unique aligned single-cell library preps. The clusters (k=3) are shown in gray, black, and red. The red vertical lines mark the read cutoff based on the 95% confidence interval of the cluster with the highest unique aligned reads. LAND = lithium assisted nucleosome depletion. xSDS = crosslinking and SDS nucleosome depletion.

**Supplementary Figure 14.**
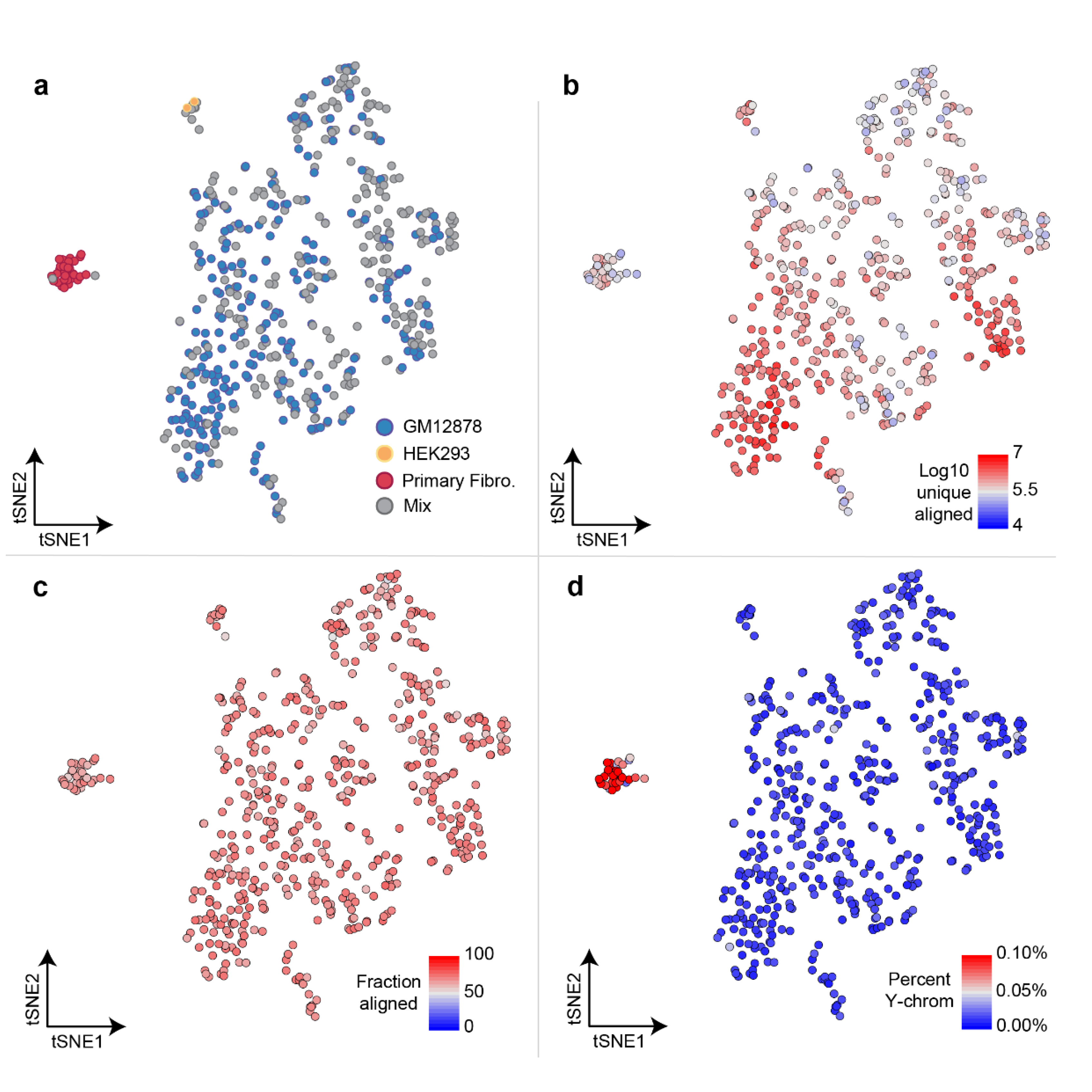
Library metrics plotted onto NMF-tSNE coordinates. a. NMF-tSNE for cells with a higher read count threshold (50,000). b. Log10 unique aligned reads with an alignment quality score ≥ 10. c. Percent of reads aligned for each library. d. Percent of reads that align to the Y-chromosome. BJ cells (foreskin fibroblast cell line) are the only cells derived from a male and form the far right cluster.

